# NIS metastable intermediates provide insights into conformational transition between principal thermodynamic states

**DOI:** 10.1101/2022.10.13.512170

**Authors:** Mayukh Chakrabarti, L. Mario Amzel, Albert Y. Lau

## Abstract

The Sodium/Iodide Symporter (NIS), a thirteen-helix transmembrane protein found in the thyroid and other tissues, transports iodide, a required constituent of thyroid hormones T3 and T4. Despite extensive experimental information and clinical data, structural details of the intermediate microstates comprising the conformational transition of NIS between its inwardly and outwardly open states remain unresolved. We present data from a combination of enhanced sampling and transition path molecular dynamics (MD) simulations that elucidate the nature of the principal intermediate states comprising the transition between the inwardly and outwardly open metastable states of fully bound and unbound NIS under an enforced ionic gradient. Our findings suggest that in both the absence and presence of bound physiological ions, NIS principally occupies a proximally inward to inwardly open state, whereas when fully bound, it also occupies a rare but thermodynamically favorable ‘inward occluded’ state. The results of this work provide novel insight into the populations of NIS intermediates and the free energy landscape comprising the conformational transition, adding to a mechanistic understanding of NIS ion transport. Moreover, the knowledge gained from this approach can serve as a basis for studies of NIS mutants to target therapeutic interventions.

For Table of Contents Only

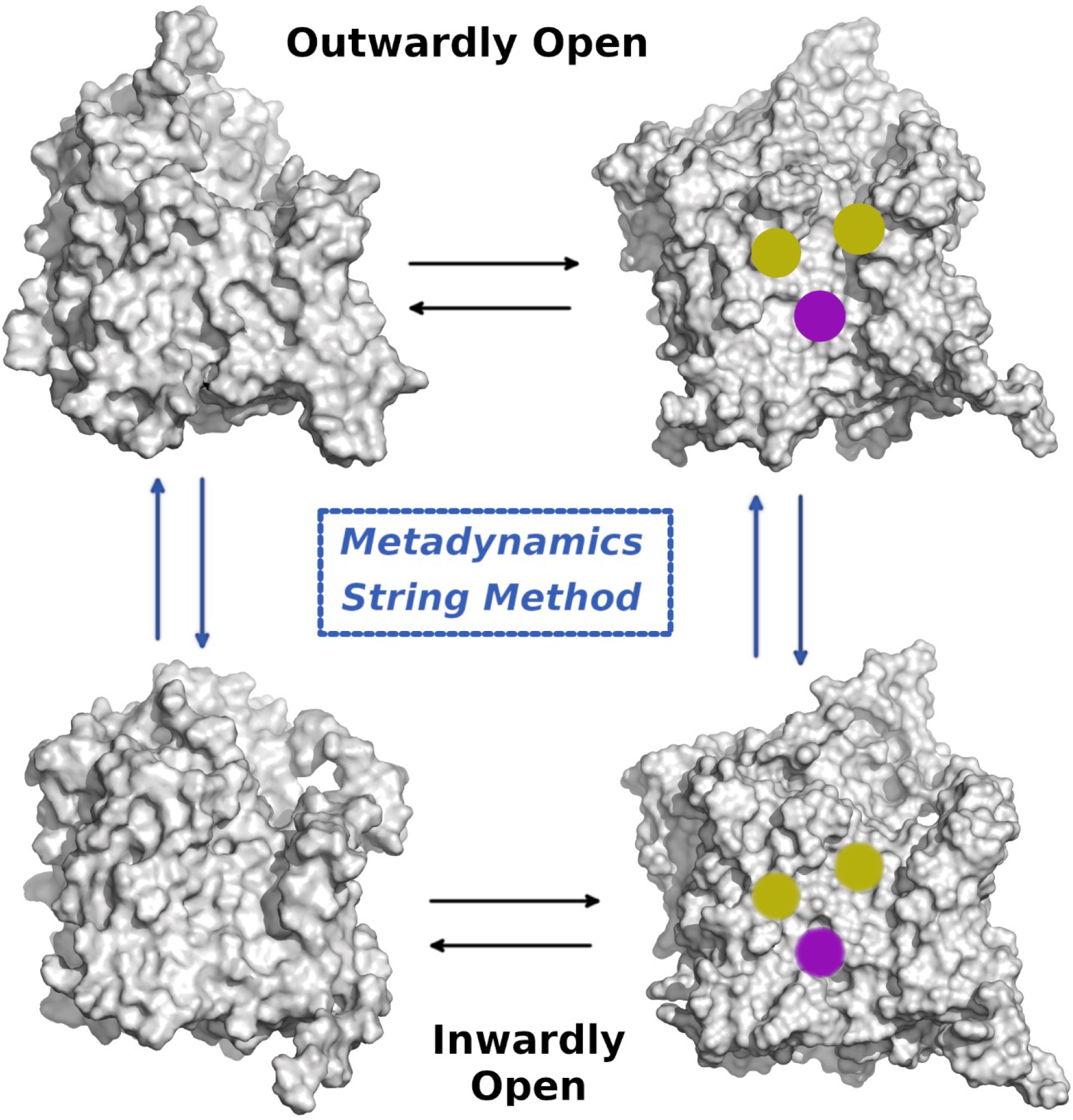

## INTRODUCTION

The Na^+^/I^−^ (NIS) symporter, a member of the solute carrier family 5 (SLCA5) Na+-dependent membrane transporters, was first characterized in the rat at the molecular level following its isolation in 1995^1^, and later isolated in humans, mice, and pigs^2–4^. It is expressed on the basolateral surface of thyroid epithelial cells, which form the follicles that comprise the thyroid gland^5^. It is also expressed in other tissues, including the salivary gland, stomach, small intestine, and lactating breast^6–8^. NIS plays a critical role in thyroid gland function and human metabolism by intracellularly accumulating iodide, which is present at sub-micromolar concentrations in the blood, allowing for the biosynthesis of the thyroid hormones triiodothyronine (T3) and tetraiodothyronine (T4)^9–11^. These hormones, in addition to regulating cellular metabolism at all stages of life in adults, play an essential role in the development of the fetal central nervous system, and are necessary for skeletal and lung growth in newborns^12^. The ability of NIS to concentrate iodide intracellularly has been utilized as one of the most effective targeted cancer therapies available: by administering radioactive iodide (^131^I^−^), thyroid cancer and hyperthyroidism can be treated ^9,11^. In humans, NIS consists of 643 amino acids, and contains three N-linked glycosylation sites that are not absolutely essential for function or stability^5,13^, but which have been shown to influence NIS trafficking and iodide uptake^14,15^. The transport of iodide by NIS is electrogenic, with a stoichiometry of 2 Na^+^: 1 I^− 16^ that provides the driving force via the downhill transport of sodium, whose electrochemical gradient across the plasma membrane is maintained by the Na^+^/K^+^ ATPase^17^. NIS is known to transport different substrates with different stoichiometries: iodide, thiocyanate (SCN^−^), and chlorate (ClO_3_^−^) are transported electrogenically with a stoichiometry of 2 Na^+^: 1 anion, whereas other oxyanions, like perrhenate (ReO_4_^−^) and perchlorate (ClO_4_^−^), are transported electroneutrally (1:1)^18^.

The current view of NIS transport involves four principal states that can be characterized in a thermodynamic cycle: outwardly open with no ions bound, outwardly open with two sodium ions and an anion bound, inwardly open with two sodium ions and an anion bound, and inwardly open with no ions bound. Until recently, the molecular structure of NIS was unknown, necessitating the development of homology models to obtain additional insights into the structural basis of NIS function. In this work, the crystal structure of *Vibrio parahaemolyticus* Na^+^/galactose transporter (vSGLT, PDB: 2XQ2), a bacterial homologue of the human Na^+^/glucose transporter SGLT1 (SLC5A1), which belongs to the same family as NIS, was used as the basis for the inward-facing homology model of rat NIS. Between core residues 50 and 456, NIS and vSGLT share 32% sequence identity and 64% similarity. For the outward-facing state of rat NIS, a homology model was generated based on the crystal structure of the *Proteus mirabilis* Na^+^-coupled sialic acid symporter (SiaT, PDB: 5NVA), which shares 25% sequence identity and 46% similarity with NIS. In the homology models, NIS consists of thirteen transmembrane helices, with an extracellular N-terminus and intracellular C-terminus, and an internal inverted repeat topology of 5-helix bundles (helices II-VI comprising repeat 1, and helices VII – XI comprising repeat 2)^10^. The inwardly open model is consistent with recently determined structures of inwardly open apo, fully bound, and perrhenate-bound rat NIS^19^. To our knowledge, no structures of outwardly open NIS have yet been determined.

The clinical manifestation of patients exhibiting iodide transport defect (ITD) disorders^20^ have led to molecular insights into the determinants of ion transport by NIS. The β-OH group of T354 in helix IX has been identified as essential for sodium binding or translocation during transport^21^, and G93 in helix III is a critical residue that is believed to facilitate the conformational change of NIS from outwardly open to inwardly open, and regulate the stoichiometry of anion transport for perrhenate and perchlorate^22^. A conformational transition mechanism that involves a 25° – 30° rotation of the helix II-helix III helical hairpin about the contact between the Cα hydrogen of residue G93 and the indole ring of W255 has been proposed based on structural modeling^22^. Mutations of G93 to T, N, Q, D, and E result in the electrogenic (2:1) transport of perchlorate and perrhenate, which are transported electroneutrally in wild-type NIS. Furthermore, G93E and G93Q mutants are unable to transport iodide at physiological concentrations^22^. Recent work has established that Y348 is a residue that influences both NIS activity and trafficking to the plasma membrane, demonstrating that charged or hydrophilic residues at this position render the protein inactive and prevent plasma membrane localization^23^. Other residues, e.g., V59, G395, R124, G543, Q267, and V270, have also been identified for their effects on NIS plasma membrane targeting and/or activity^11^.

All-atom MD simulation studies of human NIS have been previously conducted to investigate the free energy of iodide binding^24^, and to identify a putative ion translocation and gating pathway based on an inward-facing model^25^. A combination of structural data and MD simulations^26–28^ have also elucidated most of the details of the transport mechanism of *Aquifex aeolicus* Na^+^-dependent leucine transporter (LeuT)^29^, which shares the same fold as vSGLT. However, atomistic details pertaining to the conformational transition of NIS between its inwardly and outwardly open thermodynamic states in the presence and absence of bound ions, and the intermediate microstates that comprise this transition, have remained unclear. Inferences about the conformational transition profile of NIS cannot necessarily be made based on LeuT or other transporters sharing the LeuT fold, due to differences in their transported substrates, ion-substrate stoichiometry, and structure, whose effects are not predictable^30^. The recent observation that the canonical sodium binding site in LeuT does not correspond to the site of sodium ion binding in rat NIS^19^ corroborates this assertion. The difficulty of ascertaining details about the NIS conformational transition is evident from the observation that the experimentally characterized turnover rate of NIS transport in the presence of iodide is ≥ 36 s^−1 31^, corresponding to an expected inward-to-outward transition on the millisecond timescale under an unbiased sampling approach. Nonetheless, such information would provide valuable insight into the transport mechanism of NIS, elucidating the structural dynamics that occur during NIS activity and serving as a basis for therapeutic targeting of clinically manifested NIS mutants that result in transport defects. Here, we employed a combination of well-tempered metadynamics and the string method with swarms-of-trajectories, using a unique double-bilayer membrane system that permits a concentration gradient to be established and maintained across the central bilayer, to identify the populations of NIS intermediate microstates and the free energy landscape comprising the transition between inwardly and outwardly open NIS in the absence and presence of bound sodium and iodide ions. While many examples exist in the literature of such simulation approaches being utilized to examine the transition mechanisms of different transporters^32–36^, we are not presently aware of other studies that have used a combination of these approaches and applied them to the investigation of NIS dynamics, or of a simulation system in which the NIS transition mechanism was investigated under an enforced concentration gradient. As such, to the best of our knowledge, the work presented here represents a novel approach to the study of NIS transport and provides valuable structural insight into NIS conformational states that have not been accessible through other methods.

## METHODS

### MD Simulations of Inwardly and Outwardly Open NIS

Inwardly-open NIS (residues 11 – 501) was modeled using an existing structural template of the bacterial sodium/galactose symporter vSGLT (PDB ID: 2XQ2), to which it has 32% sequence identity and 64% similarity. Using the CHARMM-GUI web server^37^, the protein was solvated with the TIP3P water model and packed into a central POPC bilayer, with 110 lipids per leaflet, that is part of a 96 × 96 × 131 Å orthorhombic cell containing a total of ~125,000 atoms. The same web server was used to generate a DLPC bilayer that was manually added to the system, containing 146 lipids per leaflet, and whose purpose is to prevent ion gradients established across the NIS-containing membrane from becoming altered by periodic boundary conditions. In addition to the sodium and chloride ions used to neutralize the system, the system contains iodide ions and potassium ions. The resulting ionic concentration on both sides of the membrane was ~0.3 M, with an extracellular sodium concentration of ~140 mM and an intracellular sodium concentration of ~14 mM. A visualization of this setup is shown in Fig. 3.1. For simulations of the fully bound inwardly open state of NIS, two sodium ions, as well as an iodide ion, were placed at the positions determined through experimental assays and previous short molecular dynamics (MD) simulations^6^. Outwardly-open NIS (residues 11 – 501) was similarly modeled using an existing structural template of the sodium – coupled sialic acid symporter SiaT (PDB ID: 5NVA), to which it has 25% sequence identity and 46% similarity. Using the CHARMM-GUI web server, the protein was solvated with the TIP3P water model and packed into a central POPC bilayer, with 104 lipids per leaflet, that is part of a 92 × 92 × 149 Å orthorhombic cell containing ~117,300 atoms. As with the inwardly open system, a DLPC bilayer, containing 146 lipids per leaflet, was also present. Sodium, chloride, iodide, and potassium ions were present as in the inwardly open system, with ionic concentrations as previously mentioned. As in the case of the inwardly open system, simulations of the fully bound outwardly open state of NIS included two sodium ions and an iodide ion. A positional restraint of 2.39 kcal/mol/Å^2^ was enforced on the bound ions. For both inwardly and outwardly open NIS systems, the system energy was subsequently minimized, and equilibrated in six sequential steps: three steps of NVT equilibration with a temperature of 310K, decreasing dihedral angle restraints on the lipid molecules in the system, and decreasing z-position restraints on the lipid phosphate atoms; then, three steps of NPT equilibration at a temperature of 310K, dihedral angle restraints on the lipid molecules in the system that are reduced to zero in the final step, and z-position restraints on the lipid phosphate atoms that are reduced to zero in the final step. The protein was initially positionally restrained with a force constant of *k*_*backbone*_ = 4000 *kJ*/(*mol*) * *nm*^2^ and *k*_*sidechain*_ = 2000 *kJ*/(*mol*) * *nm*^2^, which was gradually decreased to 0 during the six sequential equilibration steps. Production MD simulations in GROMACS^38^, version 2019.4, were run in the NPT ensemble with a 2-femtosecond time-step at 310K and frames were recorded every 20 ps. All simulations used periodic boundary conditions and employed the Ewald summation methodology. A Langevin thermostat was enabled for heating the systems.

### Well-Tempered Metadynamics Simulations of Inwardly and Outwardly Open NIS

The metadynamics approach used in this work for NIS in the presence and absence of bound ions implements the procedure established by Sultan & Pande^39^. Briefly, well-tempered metadynamics was combined with supervised classification to characterize important thermodynamic states that comprise the NIS transport cycle. Unbiased sampling data was first obtained for inwardly and outwardly open NIS. For NIS with bound ions, 50 nanoseconds of simulation data were used following 600 nanoseconds and 350 nanoseconds of unbiased sampling for outwardly open and inwardly open NIS, respectively, to train a supervised classification algorithm to distinguish between the states. For NIS in the absence of bound ions, 50 nanoseconds of simulation data were used following 650 nanoseconds of unbiased sampling for both states to train a supervised classification algorithm to distinguish between the states. The training of the classification algorithm was based on protein features identified through a heuristic approach involving structural analysis and knowledge of the system. For the system with bound ions, the two sets of features used were the MET 23 Cα – SER 474 Cα distance and LYS 86 Cα – VAL 412 Cα distance. For the system without any bound ions, the two sets of features used were the THR 28 Cα – TYR 178 Cα distance and LYS 86 Cα – VAL 412 Cα distance. In both sets of simulations, the distance features are orthogonal to one another, capturing different degrees of freedom of the transporter. Using these chosen features, the classification algorithm was trained by projecting the trajectories of these features onto an XY plane and using support vector machines (SVM) to find a hyperplane that appropriately separated the two states of interest. Well-tempered metadynamics simulations were then performed using the open-source, community-developed PLUMED library^40,41^, version 2.5.3, in GROMACS 2019.4, using as a collective variable the signed distance of any sampled point to the separating hyperplane: 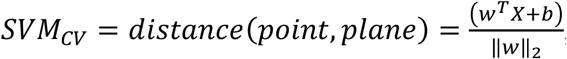, where *w* is the learned vector of collective variable coefficients, X is the input vector, and *b* is the scalar bias. Jupyter notebooks documenting the procedure, available in a public Github repository (https://github.com/msultan/SML_CV), were used to generate the SVM function used for sampling. Simulations were conducted at a temperature of 310 K, using a bias factor of 10, a gaussian kernel width (σ) of 0.1, a height of 1.0 kT, and a deposition stride of 1 ps. The sampling time for the simulation without any bound ions was 1.65 μs, and the sampling time for the simulation with bound ions was 2.1 μs.

### String Method Simulations of NIS

String method simulations were performed using a GROMACS 2021 implementation and associated Jupyter notebooks developed by the Delemotte group involving *gmxapi*^42^. For NIS in the absence of ions, the endpoint conformations for the inwardly and outwardly open states were obtained from snapshots after six hundred and fifty nanoseconds of unbiased sampling. As there was no prior knowledge about the path sampling between these two states, a linear interpolation was performed between the two endpoints, projected on an XY plane in which the abscissa was the distance between MET 23 Cα - SER 474 Cα in nm and the ordinate was the distance between LYS 86 Cα - VAL 412 Cα in nm. As noted, the images were projected on the same abscissa and ordinate as for the NIS system in the presence of bound ions in the metadynamics procedure, facilitating a direct comparison of the resulting pathways in the projected variable subspace. This resulted in thirty-three images for the initial string, later reduced to seventeen images. To obtain the initial images, a steered dynamics simulation was performed before the string simulations were initiated, using a force constant of 23.9 *kcal*/(*mol* * Å^2^) for the pull coordinates. For the string simulations, each iteration involves a restrained equilibration for 30 ps using a force constant of 23.9 *kcal*/(*mol* * Å^2^) for the pull coordinates, followed by unrestrained swarm simulations for 10 ps. One hundred twenty swarms are initiated from each image in the string in each iteration. Based on the implementation of the string method described by Pan *et al*.^43^, in which it was noted that there is no additional computational overhead associated with the inclusion of a large number of collective variables, the method was also performed using an objective selection metric for the description of factors that characterize the transition of NIS between its inwardly and outwardly open states. The paper of Pan *et al*.^43^ described a metric based on the difference in distances between all pairs of residues of a protein in its two metastable states, where all pairs with distances greater than 5 Å were chosen. In an analogous manner, to develop the selection of collective variables to be used for the procedure, all pairwise Cα distances between the inwardly and outwardly open reference states of NIS were compared, and all distance differences that were greater than 5 Å were chosen as an initial criterion. Additional filtering criteria were applied to this selection: all pairwise distances larger than half of the simulation box size were removed, and of the remaining pairs, only pairwise distances that satisfied the 5 Å cutoff metric across all images were retained. The final filtering criterion was the retention of only unique pairwise distances, such that each distance pair had a 1:1 correspondence, with each residue Cα never participating in more than one distance pair. This three-step filtering procedure resulted in 27 collective variables (listed in Supporting Information), each describing pairwise Cα distances, for NIS without bound physiological ions. The initial string for the procedure was obtained from the result following 55 iterations of the string method with two collective variables. Parameters of the string method simulations are summarized in Table 1 below.

**Table 1.** Relevant parameters and details are listed for all string method simulations performed in this work. Lists of the set of 27 variables used in this work can be found in Supporting Information.

| String Method with Swarms of Trajectories |  |  |  |  |  |  |
| --- | --- | --- | --- | --- | --- | --- |
| Bound ions | Collective Variables (Distance in nm) | Number of Images | Restrained equilibration time (ps) | Unrestrained simulation time (ps) | Number of swarms per image | Number of iterations |
| None | MET 23 CA – SER 474 CA, LYS 86 CA – VAL 412 CA | 33 | 30 | 10 | 120 | 147 |
| None | 27 variables | 17 | 30 | 10 | 120 | 43 |

## RESULTS AND DISCUSSION

### Well-Tempered Metadynamics Simulations of NIS

Well-tempered metadynamics simulations of NIS were conducted in the absence and presence of bound ions, with the objective of characterizing the conformational transition of NIS from its inwardly open to outwardly open state, as well as the populations and free energy differences of the microstates comprising the transition pathway between these two metastable states. For NIS in the presence of bound ions, data from 3 replicate metadynamics runs initiated from the inwardly open state, totaling 2.1 μs of combined sampling time, were aggregated to obtain insight into the underlying free energy landscape.

The SVM function upon which metadynamics was conducted had the following form: *SVM*_*CV*_ = (*x* + 0.022)/1.548, where the coefficients for the MET 23 Cα – SER 474 Cα distance and LYS 86 Cα – VAL 412 Cα distance collective variables were 1.505 and −0.363, respectively. Time series plots of the combined metadynamics data confirm that both sets of collective variables were diffusively sampled throughout the course of the simulations (Fig. 1). Moreover, the full range of SVM_CV_ values describing the signed distances of points from the SVM decision plane were also sampled repeatedly during these simulations (Fig. 2).

**Fig. 1.**
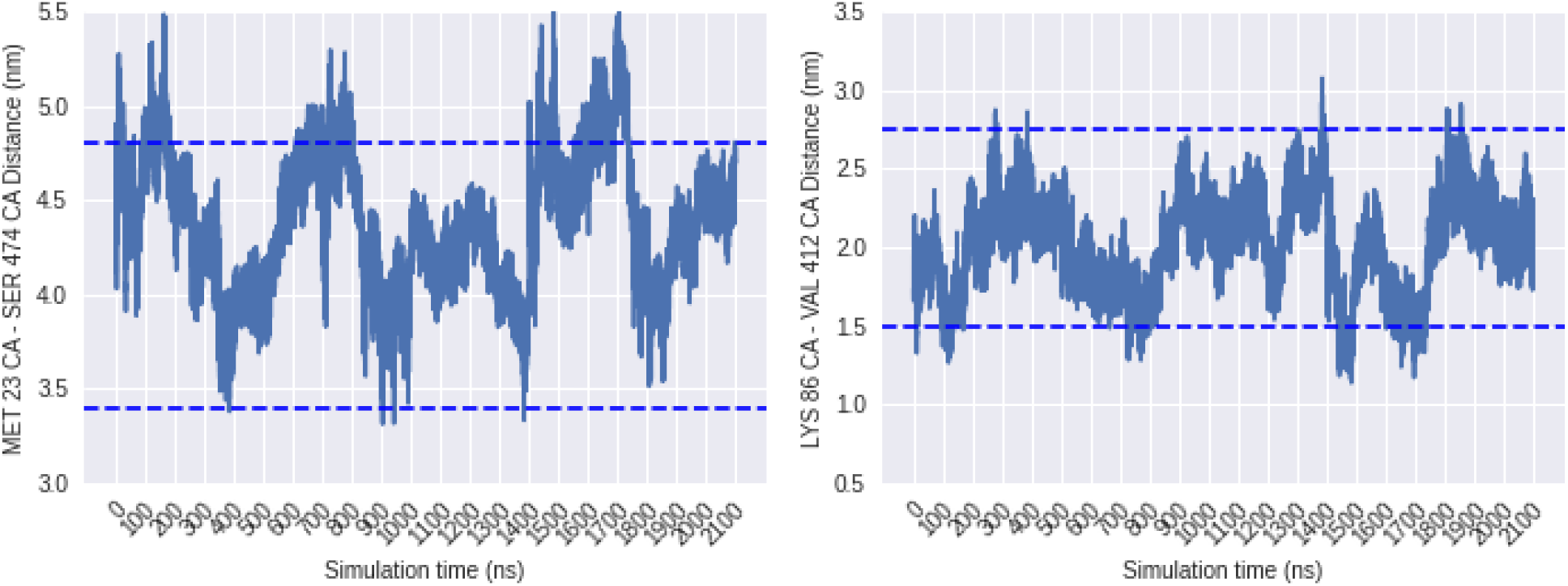
Time series plots of NIS collective variables. The values of two sets of collective variables (MET 23 CA – SER 474 CA distance, LYS 86 CA – VAL 412 CA distance) that were biased during metadynamics are shown as a function of simulation time. Data are shown as an aggregate of 3 sets of simulations with a sampling time of 2.1 μs. The blue dotted lines mark the expected sampling range of each variable.

**Fig. 2.**
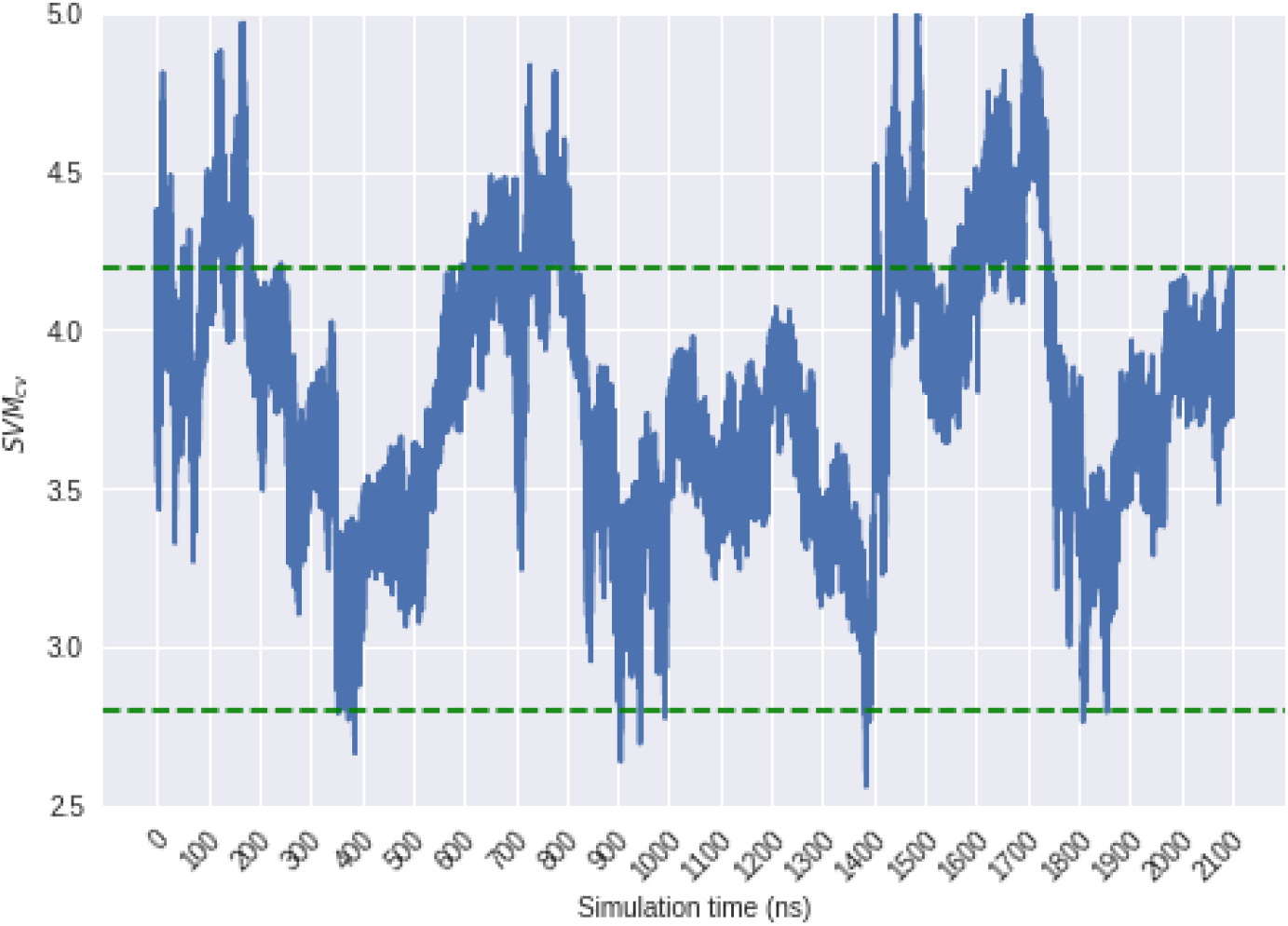
Time series plot of NIS SVM variable sampling. The value of the metadynamics collective variable, SVM_CV_, representing the signed distances from the SVM decision function that were biased during metadynamics, are shown as a function of simulation time. Data are shown as an aggregate of 3 sets of simulations with a sampling time of 2.1 μs. The green dotted lines mark the expected sampling range of the variable.

By projecting the trajectories of the collective variables onto an XY plane and performing Boltzmann inversion to obtain equilibrium ensemble weights of each observation, a free energy landscape was obtained (Fig. 3). The most relevant transition for fully bound NIS is that from the outwardly open to the inwardly open conformation, as it represents a key thermodynamic step in the transport cycle. The energy landscape suggests that NIS, in the presence of bound ions, primarily occupies two conformational states during its outwardly to inwardly open transition: an outwardly open state (labeled as State 2), and a proximally inward state that includes the inwardly open conformational ensemble (labeled as State 1). The procedure has also identified a third conformational state that can be classified as ‘inward occluded’ (labeled as State 0), representing a population that is even more closed on the outwardly facing side than the reference inwardly open structure of NIS.

**Fig. 3.**
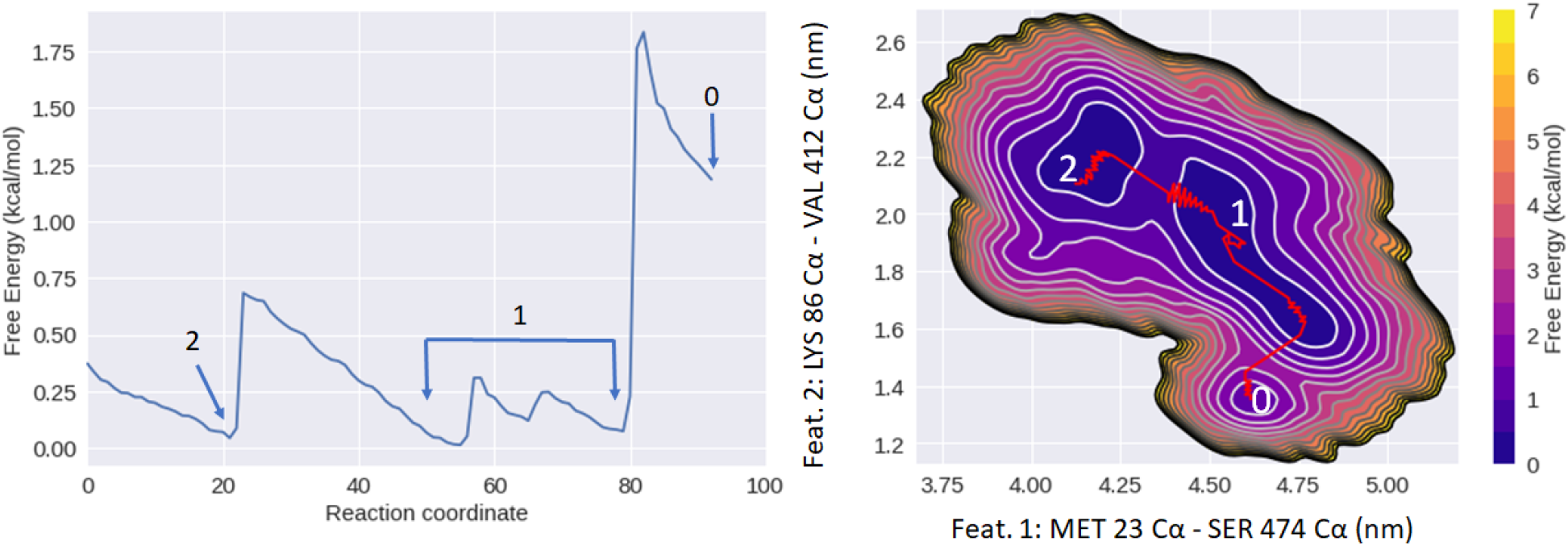
Free energy landscape of fully bound NIS from metadynamics. **Left**: A 2D projection of the minimal free energy path between the intermediates is shown as a function of a transition reaction coordinate. The identified conformational states are labeled numerically. **Right:** A 2D contour plot showing the free energy landscape projected onto the subspace of the two collective variables used in the metadynamics procedure. A minimal free energy pathway between the identified conformational states is marked in red. State labels correspond to the conformational states as marked in the 2D projection.

The significance of this ‘occluded’ population remains unclear. It could potentially represent the true inwardly open conformation of NIS, since the metadynamics simulations were performed under an imposed ionic gradient, whereas the inwardly open homology model was built from a crystal structure in which that condition was not present. Alternatively, the ensemble could potentially represent a highly populated state that occurs prior to the onset of ion release in inwardly open NIS. The minimal energy barrier separating the outwardly open state (State 2) from the inwardly open state (State 1) is on the order of thermal fluctuation 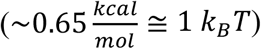. However, the barrier between the inwardly open (State 1) and ‘inwardly occluded’ state (State 0) is higher 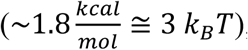, suggesting that the transition to this state may occur less frequently despite it being energetically favorable. To obtain more detailed structural insight into the nature of the three identified conformational states, representative structures were selected from each of the identified energetic minima in the landscape and examined for conformational changes. The analysis reveals numerous helical shifts and alterations in helical orientation between the outwardly open state (State 2) and the inwardly occluded state (State 0) (Fig. 4).

**Fig. 4.**
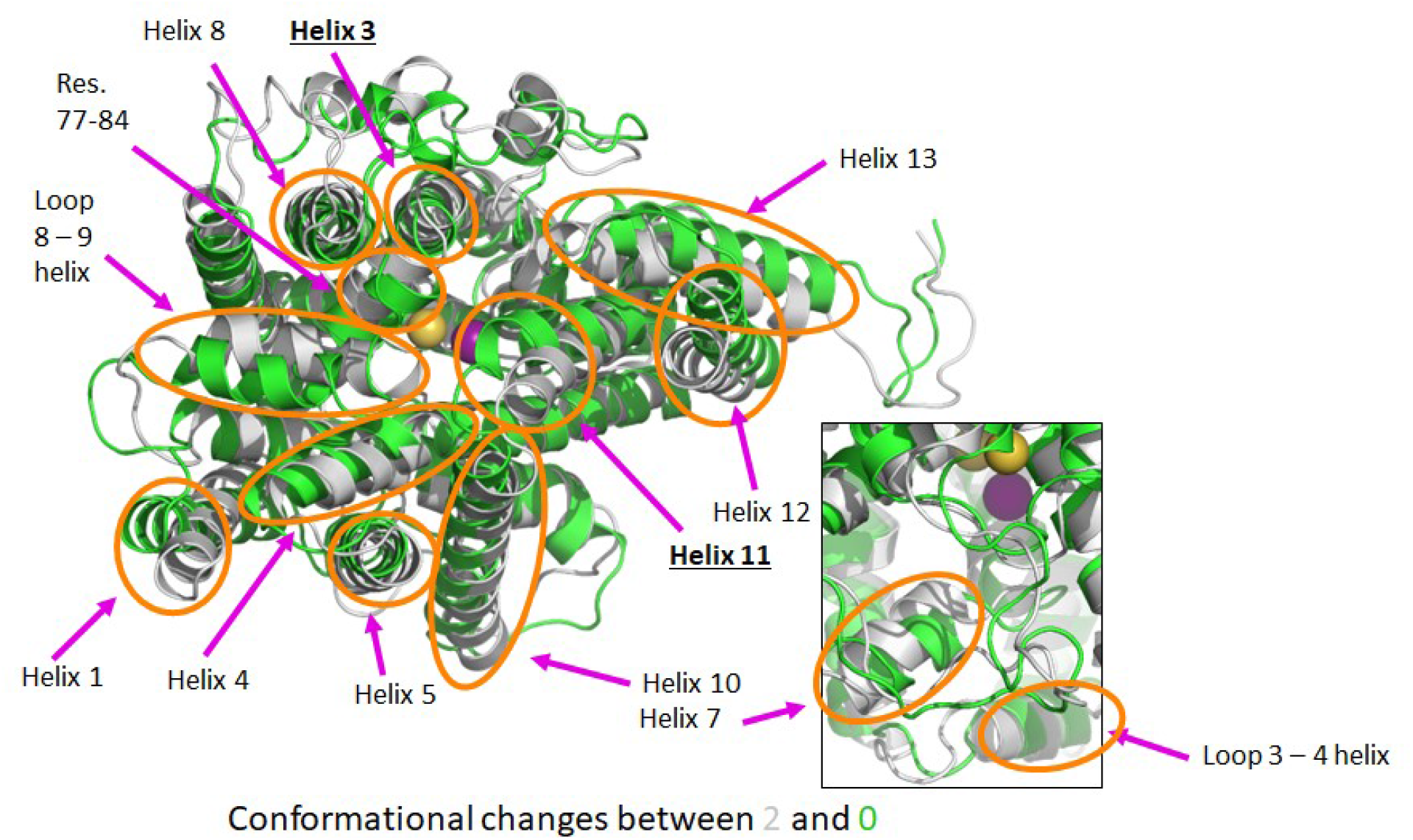
Conformational changes of NIS during its outwardly to inwardly open transition. Notable structural changes between conformational state 2 (outwardly open, gray) and state 0 (inwardly occluded, green) are shown in orange circles. NIS is shown in a bird’s eye (top-down) view, from the extracellular side of the transporter. Helices in which the changes are observed are labeled. The underlined helices include residues of the collective variable projected on the ordinate axis in the free energy landscape. In the inset figure (right), conformational changes in the inwardly facing side of NIS are marked.

The RMSD between these two states is 3.1 Å. Notably, most of these changes occur between the outwardly open state (State 2) and the proximally inward state comprising the inwardly open population (State 1) (Fig. 5), with an RMSD difference of 2.66 Å. The magnitude and extent of structural differences between the proximally inward state (State 1) and inwardly occluded state (State 0) are minimal (Fig. 6), with an RMSD of 2.0 Å. Qualitatively, this suggests that the majority of conformational changes that occur during the transition of fully bound NIS between its outwardly and inwardly open states take place during the transition between State 2 and State 1.

**Fig. 5.**
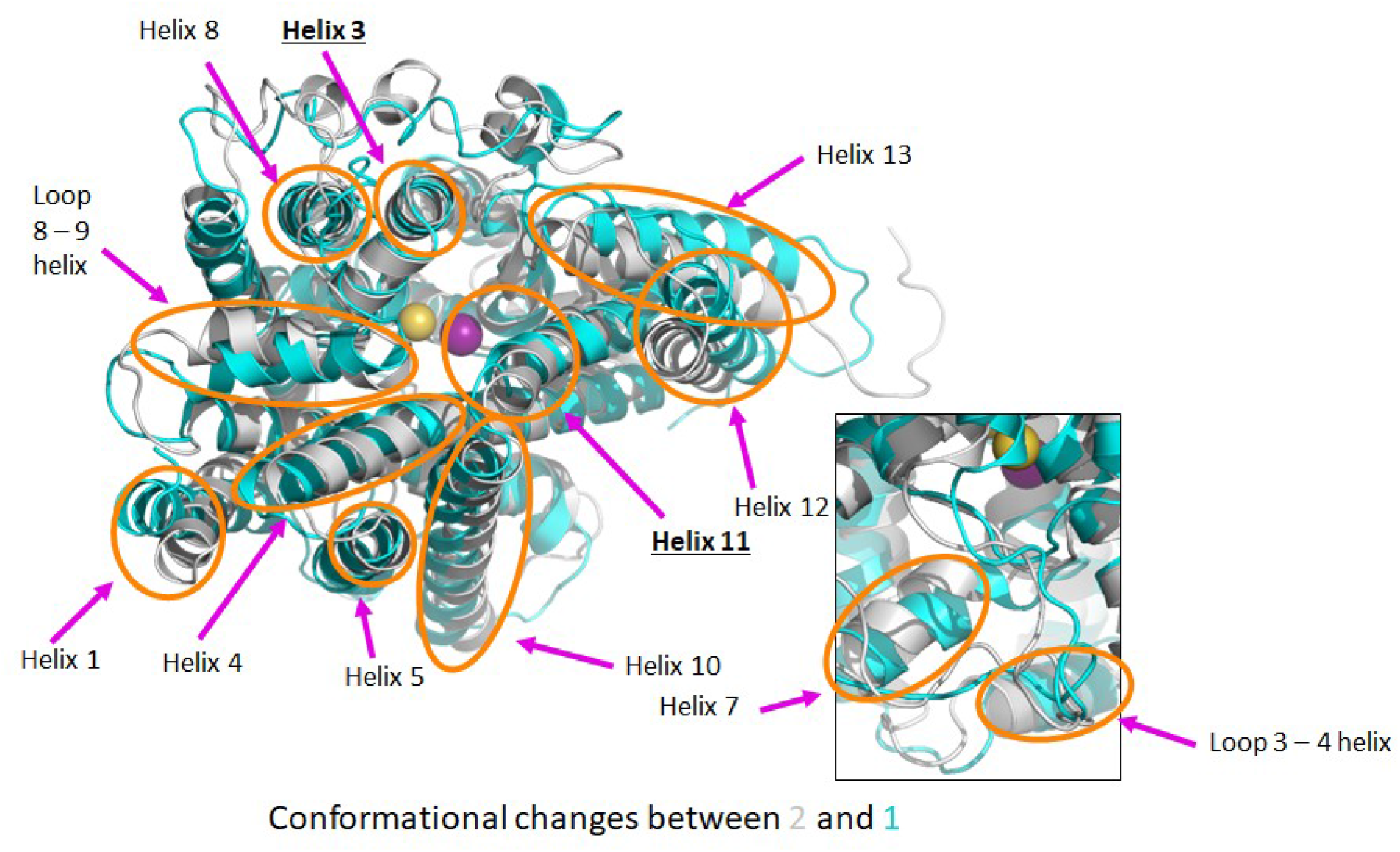
Conformational changes of NIS during its outwardly to inwardly open transition. Notable structural changes between conformational state 2 (outwardly open, gray) and state 1 (inwardly open/intermediate, aquamarine) are shown in orange circles. NIS is shown in a bird’s eye (top-down) view, from the extracellular side of the transporter. Helices in which the changes are observed are labeled. The underlined helices include residues of the collective variable projected on the ordinate axis in the free energy landscape. In the inset figure (right), conformational changes in the inwardly facing side of NIS are marked.

**Fig. 6.**
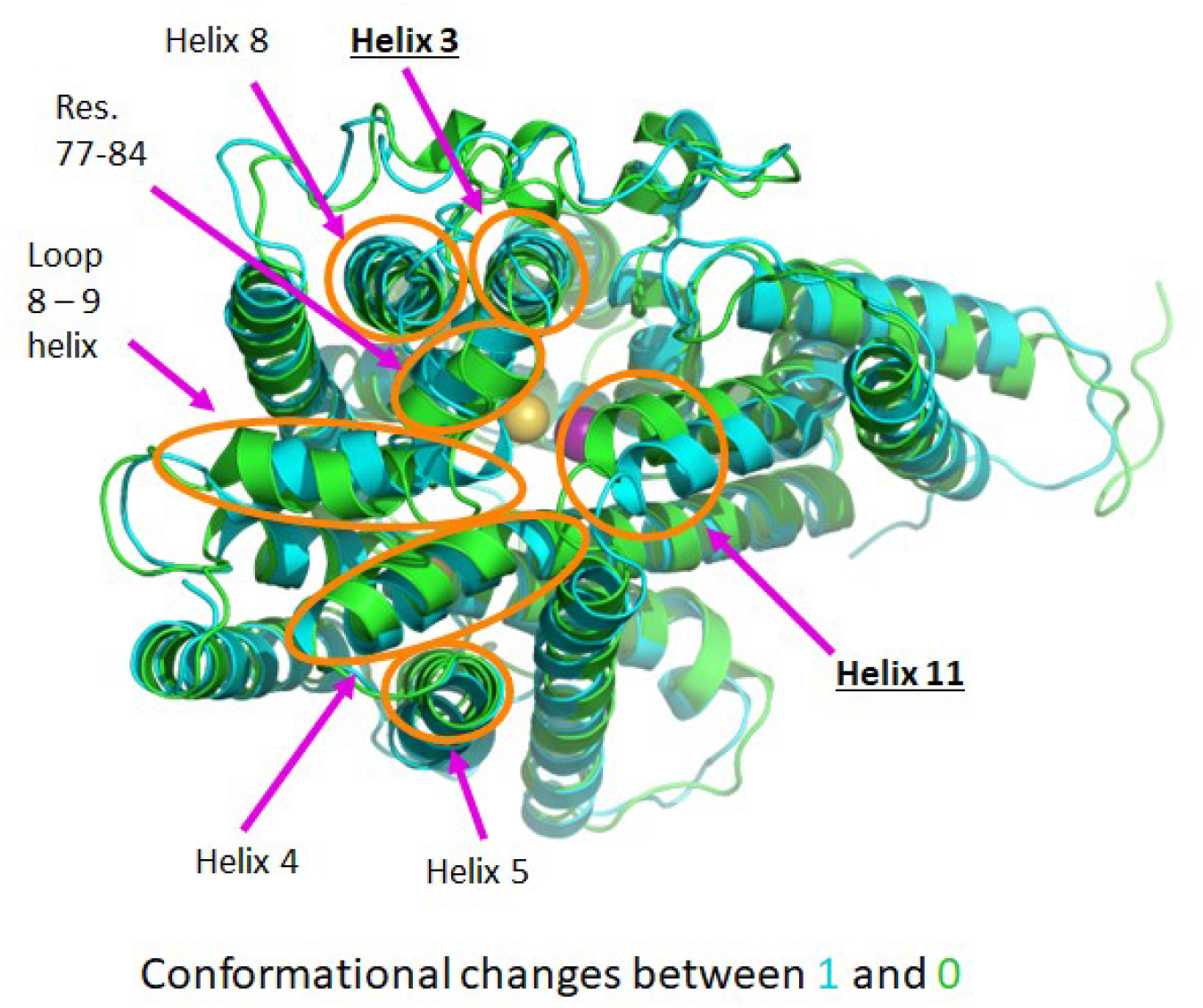
Conformational changes of NIS during its outwardly to inwardly open transition. Notable structural changes between conformational state 1 (inwardly open/intermediate, aquamarine) and state 0 (inwardly occluded, green) are shown in orange circles. NIS is shown in a bird’s eye (top-down) view, from the extracellular side of the transporter. Helices in which the changes are observed are labeled. The underlined helices include residues of the collective variable projected on the ordinate axis in the free energy landscape.

A transition path analysis, using Markov State Model transition path theory^44,45^ implemented in the Python package PyEMMA, was also conducted to examine the flux amongst the identified microstates. It reveals that the transition always proceeded via a sequential pathway from State 2 −> State 1 −> State 0, with no direct jumps between State 2 and State 0. By conducting a clustering analysis of the combined data using a mean-shift clustering algorithm (bandwidth = 0.15), four clusters were identified (Fig. 7). Importantly, the clustering yields populations of the identified conformational states. Considering cluster 1 to be the population of the outwardly open state, and clusters 0 and 2 to represent the proximally inward state that includes the inwardly open population, the populations of the states are as follows: outwardly open is 30.72%, proximally inward is 67.42%, and inwardly occluded is 1.86%.

**Fig. 7.**
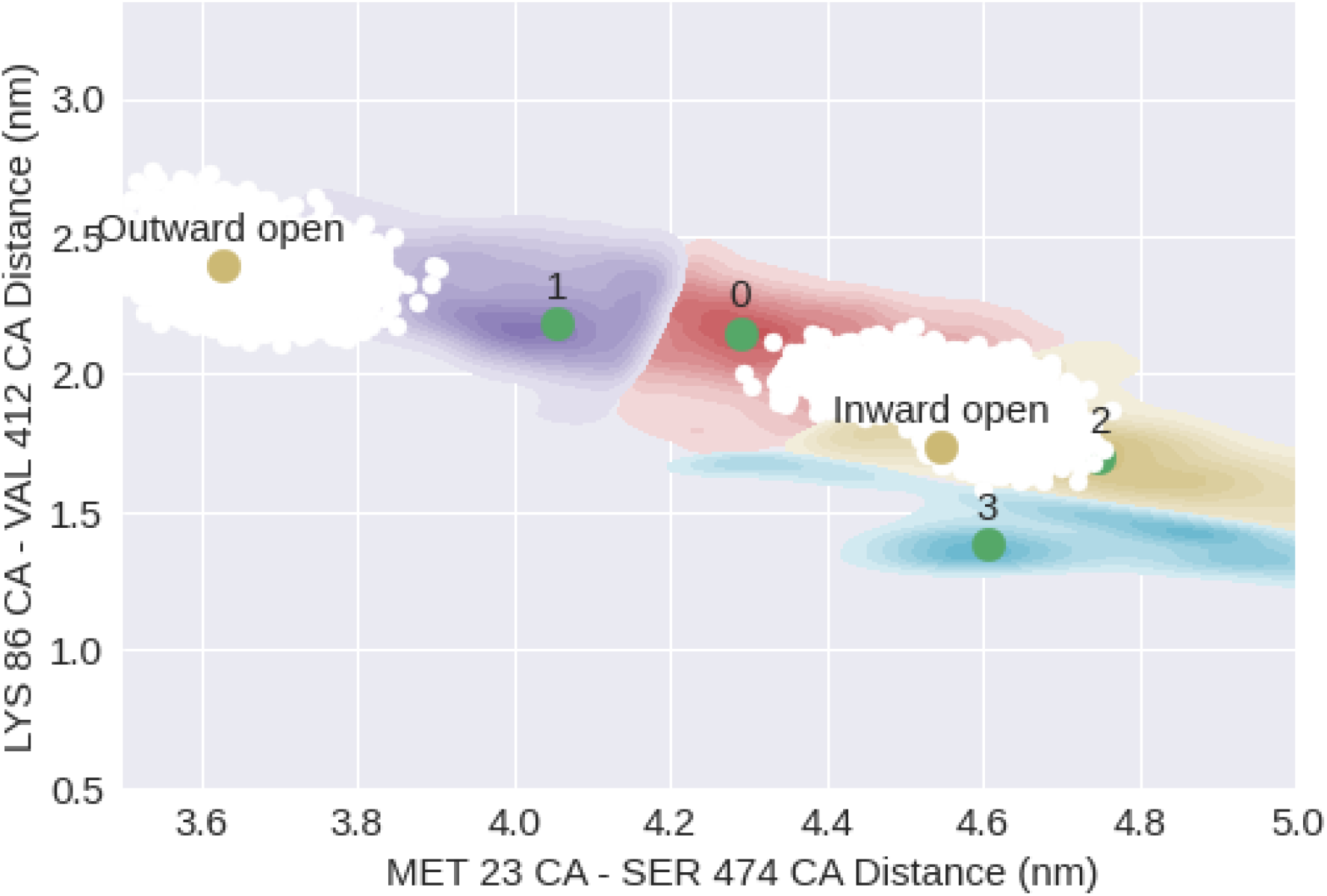
Clustering analysis of fully bound NIS. Clustering of the fully bound NIS metadynamics data using a mean-shift clustering method. Four clusters are identified. The sampling distribution of the unbiased inwardly open and outwardly open states are shown in white. Golden dots mark the location of the reference conformations used to define the inwardly and outwardly open states, as projected on the collective variable axes.

Data from three replicate metadynamics runs, totaling 1.65 μs of combined sampling time, were also obtained for NIS in the absence of bound ions. Although the inwardly and outwardly open states could be classified using SVM (*SVM*_*CV*_ = *x*/1.212, where the coefficients for the THR 28 Cα – TYR 178 Cα distance and LYS 86 Cα – VAL 412 Cα distance collective variables were 0.248 and −1.186, respectively), the procedure did not yield sampling of both metastable states, regardless of the state that was used to initiate the metadynamics procedure. Although a full range of SVM_CV_ values describing the signed distances of points from the SVM decision plane appear to have been sampled repeatedly during the simulations, time series plots of the combined metadynamics data showed that the collective variable on the abscissa (i.e., the THR 28 Cα – TYR 178 Cα distance) was not well sampled. Standard metadynamics (as opposed to the well-tempered variant) was conducted to explore the apparent kinetic barrier preventing observation of the desired transition between the inwardly and outwardly open states.

The results indicated that the transition was forced to proceed through a narrow sampling bottleneck that results in nonphysical alterations to the protein secondary structure. An attempt was made to utilize an alternative set of collective variables that would prevent structural deformations while still capturing the relevant microstates comprising the transition. Instead of a single atomic distance, the ordinate variable was changed to the center-of-mass (COM) [86 – 96 Cα] – COM [411 – 421 Cα] distance, reflecting the distance between the centers of mass of ten Cα atoms in Helix III and Helix XI, which undergo a notable change during the conformational transition between the inwardly and outwardly open states of NIS. Furthermore, the abscissa variable was changed to the MET 23 Cα – SER 474 Cα distance, the same collective variable used in the metadynamics simulations of NIS with bound ions. The SVM decision function was nearly identical to the function determined using the original set of collective variables (THR 28 Cα – TYR 178 Cα distance and LYS 86 Cα – VAL 412 Cα distance): *SVM*_*CV*_ = (*x* − 0.024)/1.199, with coefficients for the MET 23 Cα – SER 474 Cα distance and COM [86 – 96 Cα] – COM [411 – 421 Cα] distance collective variables being 0.244 and −1.174, respectively. However, this choice of variables did not resolve the barrier observed during metadynamics sampling. To obtain insight into the conformational transition of NIS in the absence of bound ions, the string method with swarms of trajectories was utilized as an alternative procedure.

### String Method Simulations of NIS

The string method with swarms of trajectories simulation approach was employed for NIS in the absence of bound ions with the objective of obtaining a minimal free energy pathway connecting the inwardly and outwardly open states of the transporter. For the string method approach using two collective variables (MET 23 Cα - SER 474 Cα distance and LYS 86 Cα - VAL 412 Cα distance), a plot of the string path by iteration for NIS in the absence of bound ions shows a significant drift relative to the original linear interpolation (Fig. 8), simultaneously demonstrating exploration of the energy landscape on the reduced collective variable subspace and providing information about the nature of the transition that was previously inaccessible using well-tempered metadynamics.

**Fig. 8.**
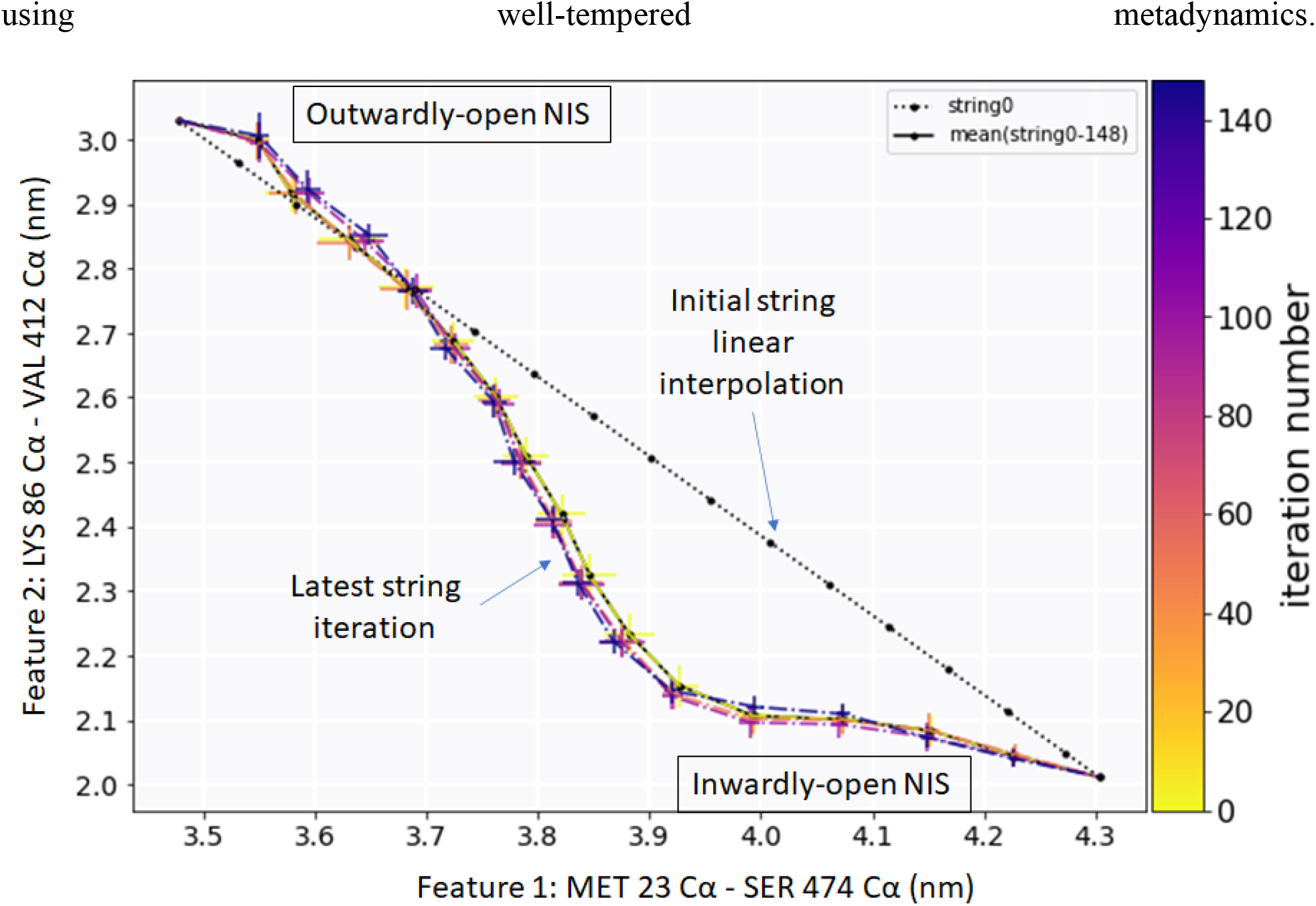
String transition pathway for NIS in the absence of bound ions. The string is projected onto a subspace of two collective variables used to describe the pathway of the transition between inwardly and outwardly open NIS. The initial string linear interpolation, used to initiate the procedure in the absence of a known transition pathway, is shown. The drift of the string by iteration number is shown by string color. The mean position and the latest position of the string are also indicated.

String convergence was evaluated using the expression 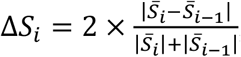, where 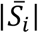 is the Euclidean norm of 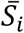, the scaled string coordinates at iteration *i*, scaled by the maximum and minimum values. The convergence plot can be found in the Supporting Information (Fig. S1). The free energy landscape in the collective variable subspace was obtained through a multi-step procedure, implemented in the *string-method-gmxapi* Python library developed by the Delemotte group and described in a recent publication^42^. Briefly, based on the two-dimensional projection of the collective variables, the procedure discretizes the system into bins, analyzes a user-defined number of swarm trajectories, and computes the number of transitions between all bins. Based on these transitions, a transition probability matrix is constructed and used to compute an equilibrium (stationary) probability distribution *p* (fulfilling detailed balance). The free energy of a given bin, *i*, is then computed using the Boltzmann expression, Δ*G*_*i*_ = −*k*_*B*_*T* log *p*_*i*_. The last 39 iterations of the string method procedure were used to obtain the swarm trajectory data used for the computation of the energy landscape. The result (Fig. 9) suggests that for NIS in the absence of bound ions, the outwardly open state is the most favorable and most highly populated. While no transition state is observable, a continuously increasing energy barrier (on the order of 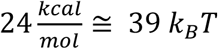 separates the inwardly and outwardly open states, suggesting that upon intracellular ion release, the transporter would quickly transition to an outwardly open state that is primed to bind sodium and iodide ions from the bloodstream.

**Fig. 9.**
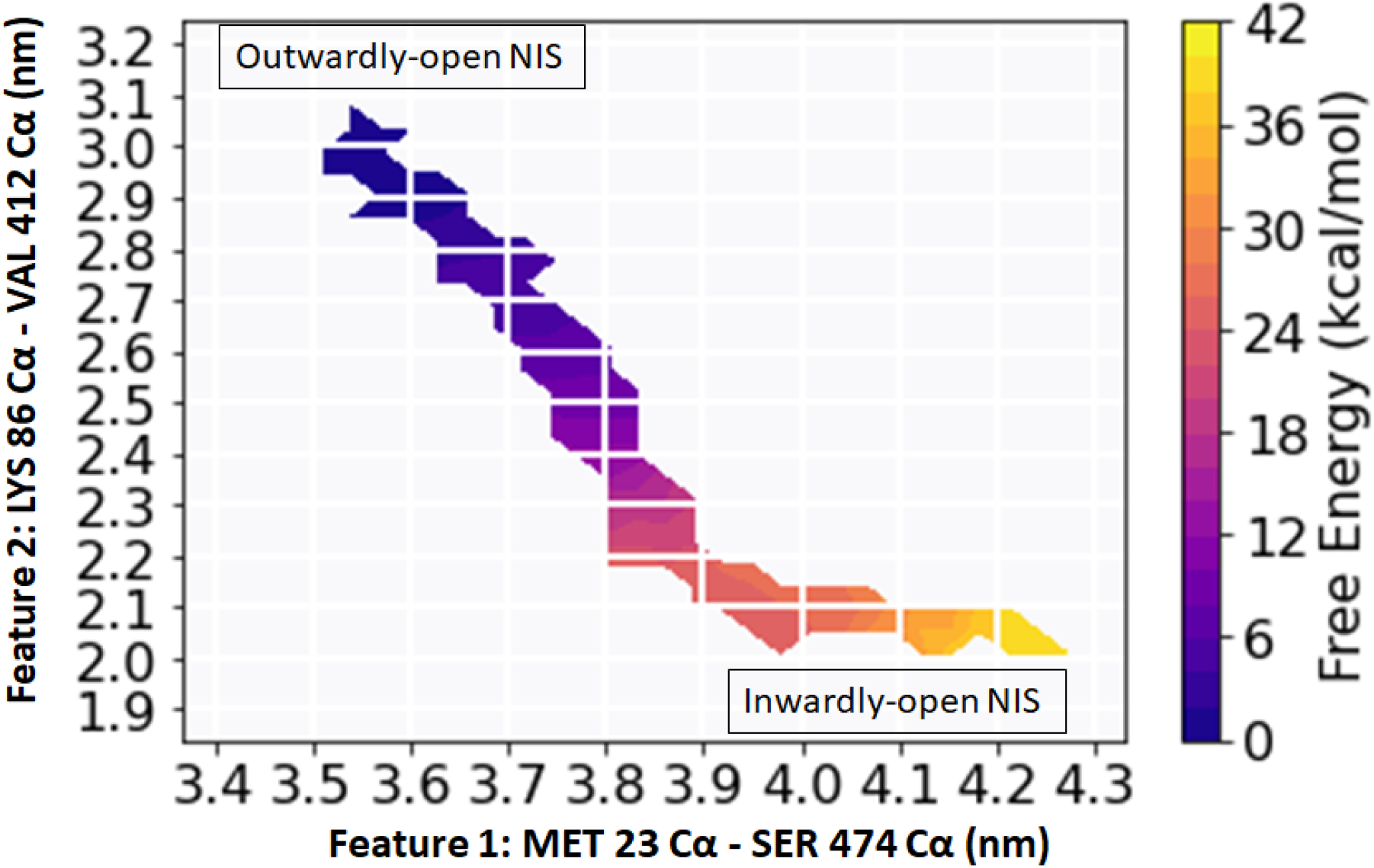
Free energy landscape for NIS in the absence of bound ions. The free energy landscape obtained by the string method procedure is projected onto a subspace of two collective variables used to describe the pathway of the transition between inwardly and outwardly open NIS.

To obtain structural insight into the microstates comprising the transition between the two metastable states, the trajectory of images comprising the string were visualized. As NIS transitions from an inwardly open state in the absence of ions, the outward movement of Helix III (containing K86) is coupled to a change in tilt of Helix X, and the change in tilt of Helix XIII (containing S474) is coupled to a change in tilt of Helix XII and a slight bend in Helix IV. These changes were also observed for the string simulations of NIS in the absence of ions conducted with 27 collective variables. To obtain insight into the free energy landscape of the conformational transition in the context of the high-dimensional space of these collective variables, the swarm trajectory data was transformed into a 1-dimensional array based on a reaction coordinate formalism and binned using the same procedure described for the two-dimensional data above. The last 34 iterations of the string procedure were used to obtain the swarm trajectory data used for the computation of the energy landscape. The reaction coordinate was defined based on the scaled Euclidean distance of the swarm trajectories from the endpoints of the string, such that values near 0 represent conformations closer to the outwardly open state, and values near 1 represent conformations closer to the inwardly open state. Interestingly, the results (Fig. 10) suggest that NIS mostly populates an ensemble of conformational states centered at a proximally inward to inwardly open state, with no evident transition state but a significant barrier to the outwardly open state on the order of 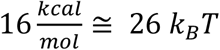. Using the same string convergence metric described previously, the data suggest that the string no longer drifts significantly in the subspace of the 27 collective variables following the fifth iteration (Fig. 11). From the underlying trajectory data, the populations of the different states can be estimated based on reaction coordinate, considering outwardly open NIS to correspond to the reaction coordinate < 0.5, a proximally inward conformation of NIS to correspond to the reaction coordinate between 0.5 – 0.7, and inwardly open NIS to correspond to the reaction coordinate > 0.7.

**Fig. 10.**
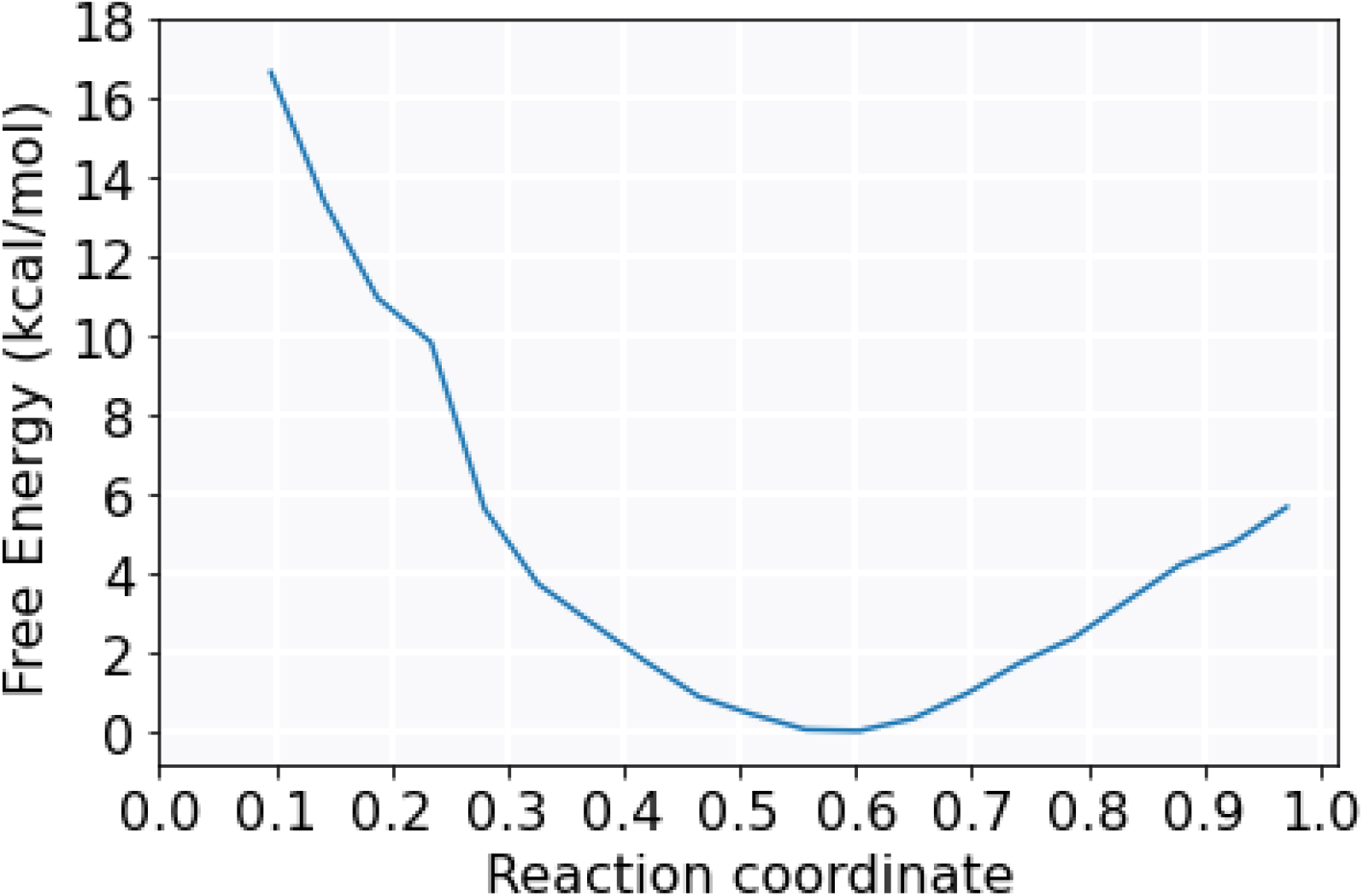
Free energy plot for NIS transition in the absence of bound ions. The free energy obtained by the string method procedure conducted on 27 collective variables is shown as a function of reaction coordinate. The coordinate is meant to describe states along the pathway of the transition between inwardly and outwardly open NIS, such that values close to 0.0 are outwardly open, and values close to 1.0 are inwardly open.

**Fig. 11.**
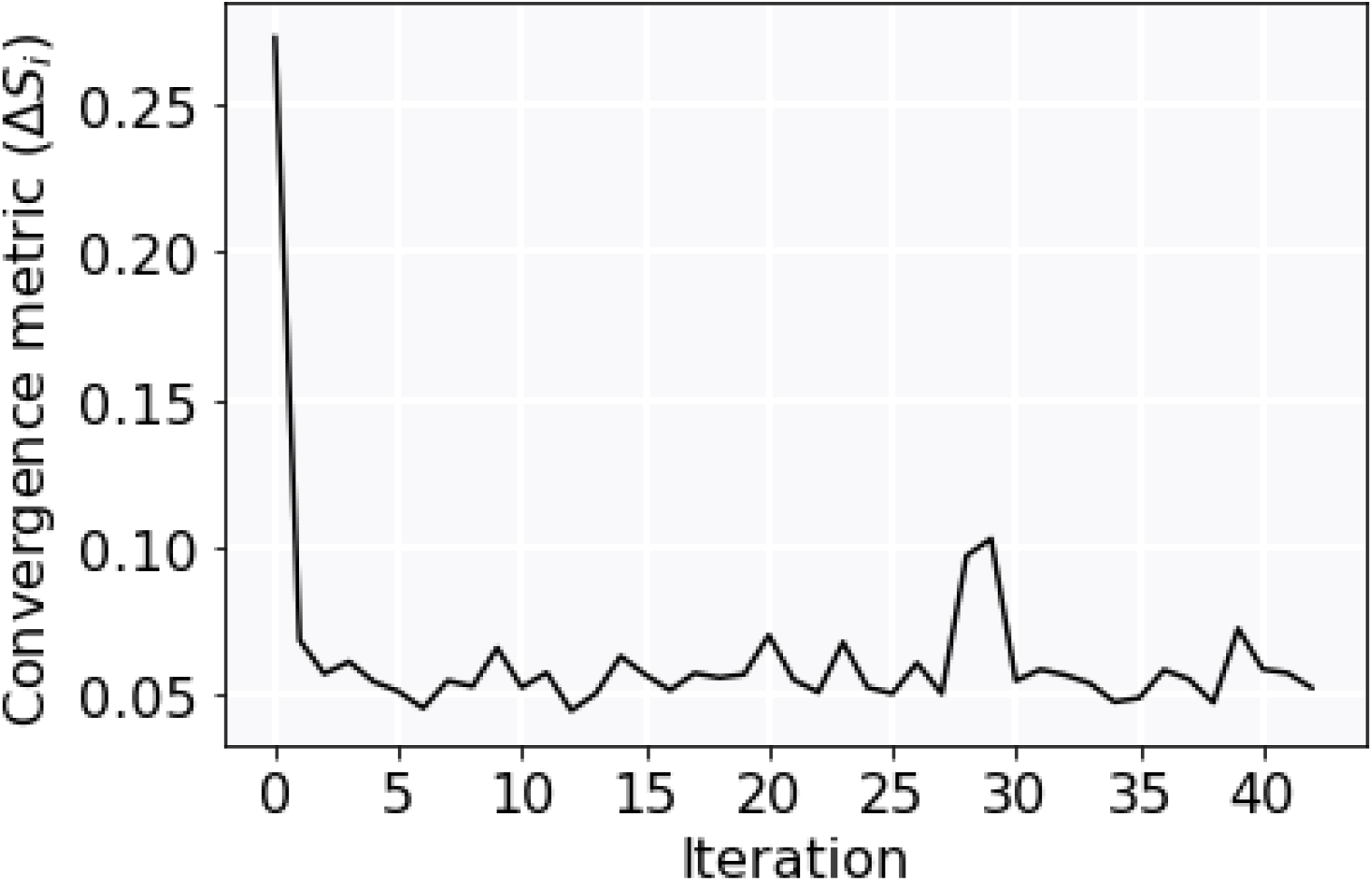
String convergence metric for NIS transition in the absence of bound ions. The string convergence metric (see text), a value used to determine the extent of string drift between iterations, is plotted as a function of iteration number.

Based on this categorization, the populations of the states are as follows: outwardly open is 36.84%, proximally inward is 34.08%, and inwardly open is 29.08%. The implication appears to be contrary to the result obtained using two collective variables: instead of the outwardly open state being highly populated, it appears that NIS in the absence of ions preferentially occupies inwardly open and proximally inward states. This could suggest that the choice of two collective variables, as previously described, may be insufficient to properly represent the conformational landscape describing the transition between the inwardly and outwardly open states of NIS in the absence of ions, and signify that the additional degrees of freedom introduced by considering several collective variables more accurately reflects the reaction coordinate of the transition, and thereby the populations of states that NIS occupies along it. The two collective variables previously described are not a subset of the string in the subspace of 27 collective variables, and thus an alternative explanation for the observed result could be the fact that the landscape does not correspond to the same reaction coordinate.

## CONCLUSIONS

A combination of well-tempered metadynamics simulations and path sampling approaches were used in the present study to characterize the conformational dynamics of NIS from its inwardly open to outwardly open state in the absence and presence of bound physiological ions, and provide novel and valuable insight into these processes. These transitions represent two critical steps in a simplified thermodynamic cycle of NIS activity, comprised of many additional thermodynamic states whose temporal order and mechanistic details remain to be fully elucidated^12,19^. Implicit in this cycle, and in the work presented, is the notion of an ‘alternating access’ model that is characteristic of transporters^46–48^, in which conformational transitions that occur between outwardly and inwardly facing states alternatively expose the substrate (in the case of NIS, a bound anion) to the extracellular or cytoplasmic milieu. Although NIS was modeled based on the structure of vSGLT, which shares the same fold as LeuT, the data for fully bound NIS intriguingly does not appear to support a ‘rocking-bundle’ mechanism as would be expected from a transporter of this fold^46,49^. Helices II, III, VII, and VIII of NIS have been previously noted to comprise the bundle domain, and helices IV, VI, IX, and XI have been noted to comprise the scaffold domain^10^. Instead of a movement of the bundle domain relative to the scaffold, it rather appears that the transition between the outwardly and inwardly open states in the presence of bound ions may be accompanied by distinctive movements in peripheral helices I, XII, and XIII, with the position of bundle helix III changing and helix XI of the scaffold also shifting during the outwardly to inwardly open transition (Fig. S2).

In the absence of bound ions, NIS likely predominantly occupies a proximally inward to inwardly open state. Physiologically, it could be surmised that the binding of a sodium ion to this state would drive NIS towards an outwardly open state and prime it for a subsequent cycle of ion transport. Observations of the conformational dynamics of other sodium-binding transporters have suggested this to be a plausible hypothesis^34,50–52^. The findings presented here suggest that the ‘futile cycle’ of NIS transitioning from an outwardly to inwardly open state in the absence of bound ions may involve an inward movement of Helix III, together with changes to the helical tilt of Helices X, XII, and XIII, and a helical bend in Helix IV. When fully bound with physiological ions, the data suggest that NIS occupies two metastable states: an outwardly open state and a broad proximally inward state that includes the inwardly open ensemble. The population of fully bound NIS that is proximally inward to inwardly open appears to be greater than that of the outwardly open state, suggestive of the fact that NIS is more likely to be poised to release its ions into the intracellular milieu rather than remain in an outwardly open state. Taken together, the findings presented in this work are a step towards addressing many of the current questions in the field about the mechanistic details of NIS transport activity and the characterization of its conformational states, and serve as a basis for future experimentally testable hypotheses.

## SUPPORTING INFORMATION

Table S1. List of 27 collective variables (CVs) used for the string method simulation in the absence of bound ions.

Figure S1. Plot of the string convergence metric for NIS in the absence of bound physiological ions, conducted using 2 collective variables.

Figure S2. Transition of NIS between the outwardly open and inwardly occluded states in the presence of bound ions, with ‘bundle’ and ‘scaffold’ domains labeled.

## AUTHOR INFORMATION

### Author Contributions

The manuscript was written through contributions of all authors. M.C. was involved in Conceptualization, Methodology, Software, Validation, Formal Analysis, Investigation, Writing – Original Draft, and Visualization. L.M.A. was involved in Conceptualization, Methodology, Resources, Visualization, Supervision, Project Administration, and Funding Acquisition. A.Y.L. was involved in Supervision, Visualization, Writing – Review & Editing, and Validation. The authors have given approval to the final version of the manuscript.

## ACKNOWLEDGMENTS

We thank Dr. Mario A. Bianchet for helpful advice and discussions. We used CPU and GPU resources and scientific computing services at the Maryland Advanced Research Computing Center (MARCC) and Advanced Research Computing at Hopkins (ARCH), both at Johns Hopkins University.

## ABBREVIATIONS

NIS: Sodium/Iodide Symporter
MD: molecular dynamics
SLCA5: solute carrier family 5
T3: triiodothyronine
T4: tetraiodothyronine
SCN^−^: thiocyanate
ClO_3_^−^: chlorate
ReO_4_^−^: perrhenate
ClO4^−^: perchlorate
vSGLT: *Vibrio parahaemolyticus* Na^+^/galactose transporter
SiaT: *Proteus mirabilis* Na^+^- coupled sialic acid symporter
ITD: iodide transport defect
LeuT: *Aquifex aeolicus* Na^+^-dependent leucine transporter
POPC: 1-palmitoyl-2-oleoyl-sn-glycero-3-phosphocholine
DLPC: 1,2-dilauroyl-sn-glycero-3-phosphocholine
SVM: Support Vector Machine

