## Supplemental Information for "NIS metastable intermediates provide insights into conformational transition between principal thermodynamic states"

[#]L. Mario Amzel passed away on August 28, 2021.

| | | | |
|---|---|---|---|
| THR 11 CA – LEU 314 CA | LEU 24 CA – LEU 313 CA | GLY 51 CA – ALA 439 CA | ALA 56 CA – VAL 412 CA |
| VAL 57 CA – GLY 411 CA | MET 68 CA – SER 466 CA | SER 69 CA – TYR 471 CA | ALA 70 CA – MET 467 CA |
| VAL 71 CA – PRO 464 CA | ARG 82 CA – TYR 477 CA | TYR 83 CA – LEU 408 CA | GLY 84 CA – GLY 410 CA |
| LEU 85 CA – LEU 413 CA | PRO 107 CA – LEU 378 CA | ILE 108 CA – CYS 440 CA | ARG 111 CA – GLY 377 CA |
| LEU 112 CA – ALA 379 CA | GLY 113 CA – PRO 380 CA | SER 116 CA – HIS 273 CA | VAL 139 CA – ARG 317 CA |
| THR 160 CA – PRO 337 CA | THR 171 CA – GLN 414 CA | ILE 174 CA – TYR 475 CA | HIS 226 CA – GLY 409 CA |
| LEU 231 CA – GLY 415 CA | CYS 310 CA – SER 319 CA | PRO 438 CA – PHE 462 CA | |

**Table S1. List of collective variables used in the string method of NIS without bound physiological ions.** All variables are atomic distances.

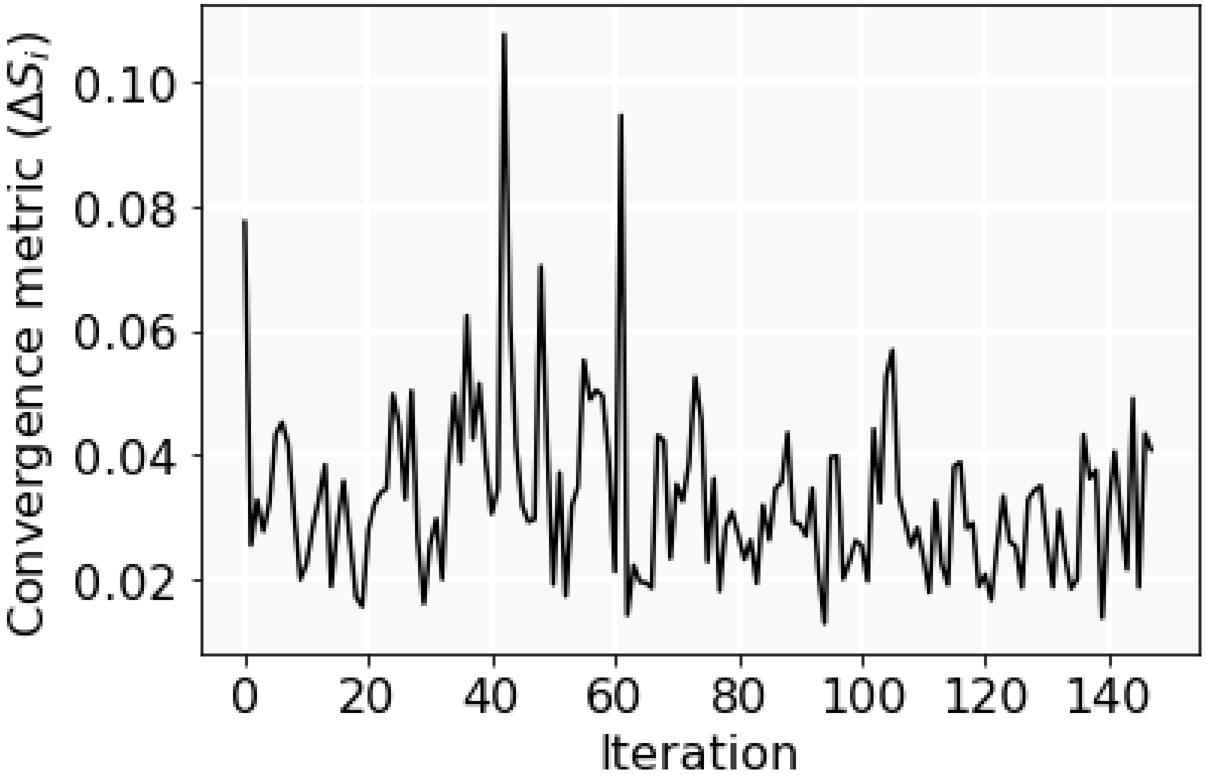


**Fig. S1. String convergence metric plot for NIS in the absence of bound physiological ions, conducted using 2 collective variables.** The string convergence metric (see main text), a value used to determine the extent of string drift between iterations, is plotted as a function of iteration number.

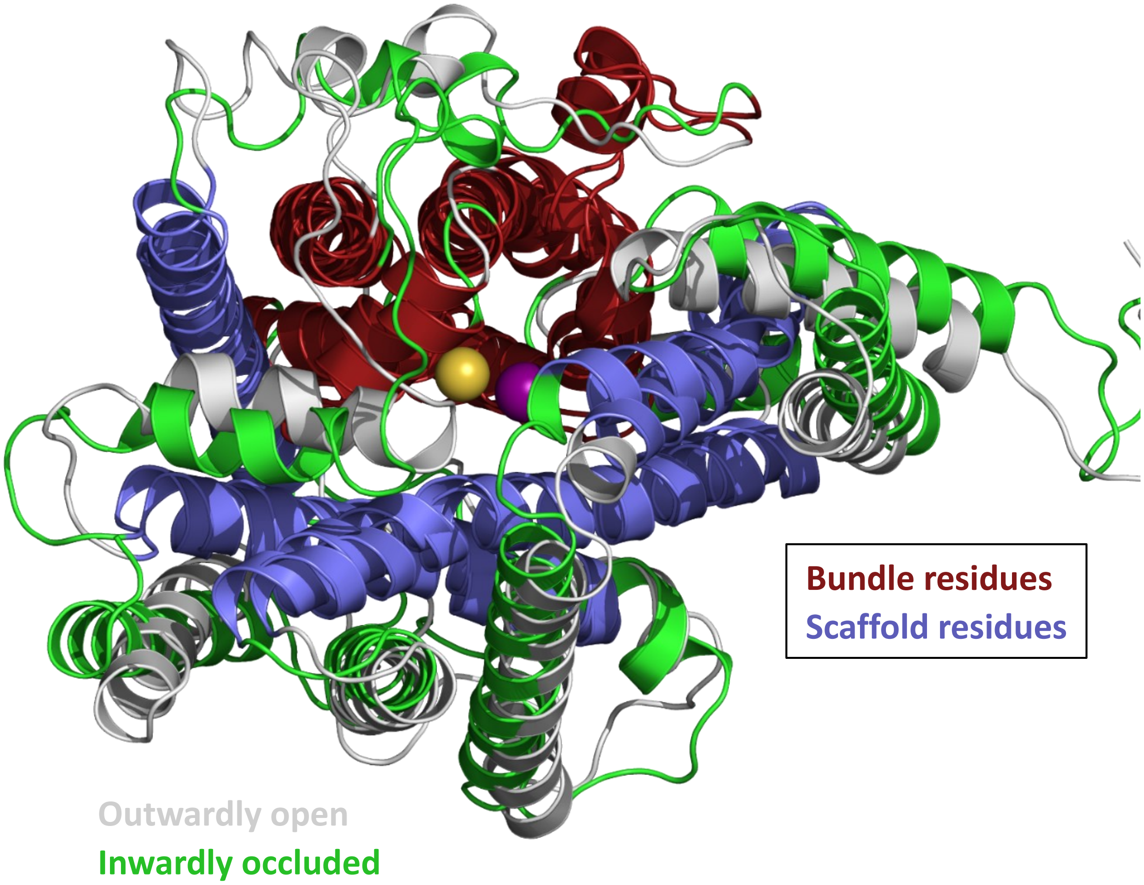


**Fig. S2. Transition of NIS between the outwardly open and inwardly occluded state in the presence of bound ions.** The 'bundle' and 'scaffold' residues of NIS, comprising Helices II, III, VII, and VIII of NIS, and helices IV, VI, IX, and XI of NIS, respectively, have been labeled. NIS is shown in a bird's eye (top-down) view, from the extracellular side of the transporter.
